# Expanding the methionine toolkit: *L*-cyanohomoalanine as a multifunctional analog

**DOI:** 10.64898/2026.06.25.734610

**Authors:** Sydney O. Shuster, Caitlin M. Davis

## Abstract

Non-canonical amino acids (ncAAs) are valuable tools in chemical biology and biochemistry for labeling, probing, and tracking biomolecules. ncAAs that can be recombinantly incorporated using native *E. coli* machinery are particularly useful because they allow for global protein incorporation and avoid complex genetic code expansion. Here, we demonstrate successful incorporation of a methionine analog, *L*-cyanohomoalanine (Cha), by the methionyl-tRNA synthetase of *E. coli* into mutant superfolder GFP (sfGFP) expressed in methionine auxotroph bacterial cultures. We compare to methionine auxotroph bacterial cultures supplemented with *L-*methionine (Met) or *L*-azidohomoalanine (Aha). In control prototrophic *E. coli*, bacterial growth rates are inhibited with high concentrations of Aha but not Cha. However, less sfGFP is produced in auxotrophic cells supplemented with Cha compared to Aha and Met. Thus, while Cha is non-toxic to *E. coli* it is incorporated less efficiently into proteins than Aha or Met. Mass spectrometry confirmed that N-terminal Cha, Aha, and Met are cleaved, as expected for the sfGFP mutants. Other sites of Cha and Aha incorporation were confirmed by mass spectrometry, with labeling efficiency varying by position. Thermal melts of purified sfGFPs demonstrate that Cha and Aha labeling does not significantly perturb the protein stability. In the future, Cha may be useful for proteome labeling by wild-type methionyl-tRNA synthetase and could be implemented in metabolic pulse-labeling of newly synthesized proteins with other methionine analogs. Additionally, the nitrile moiety of Cha may be used to perform reactions orthogonal to azide/alkyne click chemistry or could serve as a vibrational reporter of the environment.

## Introduction

Noncanonical amino acids (ncAAs) are valuable research tools used to track protein localization within the cell, characterize protein structure and local environment, and serve as a handle for bioorthogonal chemistry.^1–3^ Amber codon suppression and genetic code expansion (GCE) have allowed for the introduction of a near limitless class of ncAAs.^4,5^ However, GCE can be difficult to implement and is only desirable for site specific incorporation. Alternatively, ncAAs can be incorporated by making use of the natural promiscuity of the methionyl-tRNA synthetase (metRS) to globally replace *L-*methionine (Met) residues, most notably with *L-*azidohomoalanine (Aha),^6^ *L-*homopropargylglycine (Hpg),^7^ and selenomethionine (SeMet).^8^ Here, we describe the metRS mediated incorporation of another ncAA, (*S*)-2-Amino-4-cyanobutanoic acid, into a protein of interest and compare its incorporation to Aha. Following the naming convention of Aha, we refer to (*S*)-2-Amino-4-cyanobutanoic acid as *L-*cyanohomoalanine (Cha) as it contains a nitrile group (cyano-) and is a higher homolog (homo-), containing one additional methyl group, of alanine.

MetRS promiscuity is well documented and has been previously exploited to incorporate methionine analogs with azide and alkyne functionalization (Aha and Hpg, respectively) as well as photoreactive *L-*photo-methionine (pMet) and SeMet^6,7,9,10^ Both Aha and Hpg were originally developed as click chemistry handles for copper-catalyzed azide-alkyne cycloaddition (CuAAC) reactions but are also useful probes for vibrational spectroscopy.^11,12^ Azide and alkyne moieties absorb in the “cell-silent” region (1900–2400 cm⁻¹) of infrared (IR) and Raman spectra, respectively, and are sensitive to local environment, providing information on both the amino acid location and microenvironment.^1,13^ However, each probe has limitations. For example, Aha can induce cellular stress, restrict growth, and alter protein expression.^14–16^ Alkyne moieties can also affect cell growth^16^ and, despite being excellent Raman probes, exhibit weak IR absorptions due to the symmetric nature of the alkyne stretch and thus cannot serve as IR probes.^17^

Nitrile moieties offer a complementary set of capabilities that could expand the Met analog toolkit. Because nitrile moieties possess intermediate polarity between amide and methylene moieties, it may be possible to more flexibly incorporate them into protein hydrophobic cores and hydrophilic surfaces than alkyne moieties.^18–21^ Nitriles are also capable of versatile chemistry: Cha has already been used in peptide synthesis as a glutamine synthon^22^ and nitrile containing drugs have been used to covalently crosslink with cysteine residues.^23^ Click chemistry of nitrile and allene moieties has recently emerged as an orthogonal clickable pair to azide and alkyne moieties, creating a need for new ncAA systems compatible with this chemistry.^24^ Additionally, nitriles are useful MS and vibrational probes. Like azide and alkyne moieties, nitrile moieties are visible in the cell-silent region of both Raman and IR spectroscopy. Nitrile containing amino acids, most notably *p*-cyano-*L-*phenylalanine (*p*CNPhe) ^21,25^ have already been incorporated via peptide synthesis and genetic code expansion for IR studies. Despite these advantages, no nitrile-labeled Met analog has been expressed in proteins using native cellular machinery.

In this work, we demonstrate that Cha, a nitrile containing Met analog that is natively produced by Chromobacteria,^28^ can be successfully incorporated into model proteins by methionine auxotrophic *E. coli* during methionine deprivation using metRS (Fig. 1). Incorporation levels are low, but Cha is well tolerated by *E. coli* and offers a multi-modal probe that expands the Met analog toolkit available for protein studies. Future work may evolve *E. coli* to better incorporate Cha, which will expand its utility both in bioorthogonal chemistry and as a vibrational or mass spectrometry probe.

**Figure 1.**
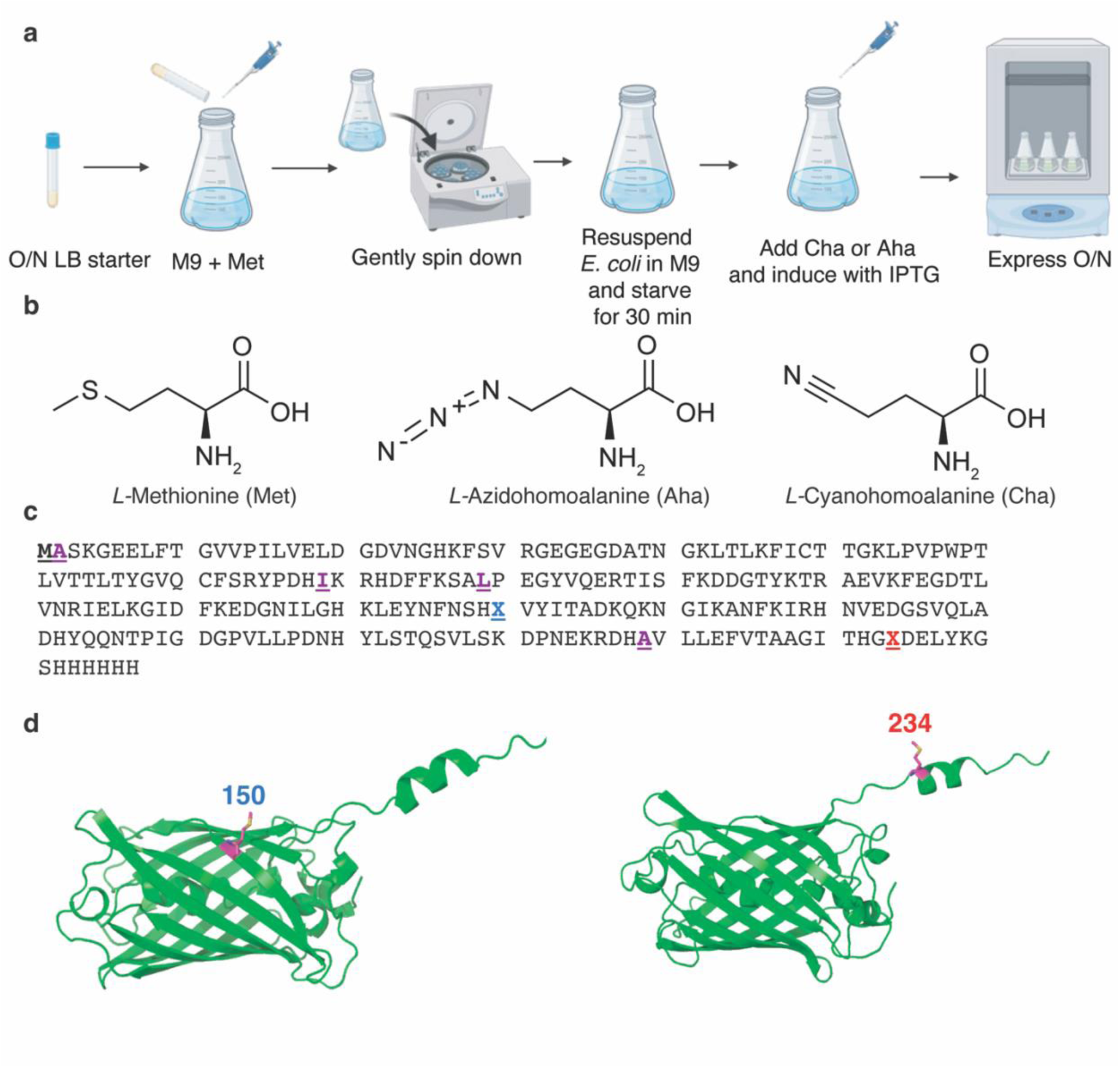
*L-*cyanohomoalanine (Cha) and *L-*azidohomoalanine (Aha) incorporation to mutant superfolder GFP (sfGFP). (a) Protocol for Cha and Aha incorporation into sfGFP. Methionine auxotroph *E. coli* transformed with sfGFP are grown in 5 mL of Luria broth (LB) overnight (O/N). This starter is transferred to 1 L of M9 media (minimal media, supplemented with 50 mg/L *L*-methionine (Met)). The *E. coli* is grown at 37 °C until it reaches an optical density (OD) at 600 nm of 0.8-1 and then collected via centrifugation. The *E. coli* are resuspended in fresh M9 media and starved at 37 °C for 30 minutes. Cha or Aha is supplemented (40 mg/L) and protein expression induced with IPTG (1mM). Expression occurs over 16 hours at 20 °C. Workflow created with Biorender.^26^ (b) Chemical structures of Met, Aha, and Cha. Structures generated in ChemDraw. (c) Sequence of sfGFP mutated to contain Met only at the N-termini and position 150 or 224. Residues in purple have been mutated from sfGFP in both 150 and 234 sfGFP. The blue underlined X is a Met, Aha, or Cha in 150 sfGFP and an Asp in 224 sfGFP. The red underlined X is a Met, Aha, or Cha in 234 sfGFP and an Ala in 150 sfGFP. The bold underlined N-terminal methionine is proteolytically cleaved in *E. coli*. (d) Ribbon diagrams of Alphafold 3^27^ structure predictions of 150 Met sfGFP (left) and 234 Met sfGFP (right) with the Cha/Aha incorporation position colored magenta and label colored to match (c). Ribbon diagrams generated in PyMol. Figure assembled in Adobe Illustrator.

## Results and discussion

We first assessed the effect of Cha and Aha on *E. coli* growth in M9 media alone or containing Cha or Aha at three concentrations with a Met prototroph *E. coli* line (BL21 DE3). The optical density (OD) at 600 nm of the growths was followed over 24 hours (Fig. 2, Fig. S1). Growth was assessed when cultured reached an OD of 0.8, where protein expression would typically be induced. We found that Cha did not affect *E. coli* growth at 40 or 200 μg/mL and only marginally slowed growth at 800 μg/mL (Fig. 2a). Aha did not affect *E. coli* growth at 40 μg/mL but began to slow growth at 200 μg/mL with increased effects at 800 μg/mL (Fig. 2b). This agrees with past work that found higher concentrations of Aha can cause toxicity and cell stress.^16^ Although *E. coli* tolerate more Cha than Aha, to reduce undesired side effects from toxicity (e.g. changes in post-translational modifications^14,29^) and allow for direct comparison of expression yields, moving forward we supplemented protein expressions with 40 μg/mL for both Aha and Cha.

**Figure 2.**
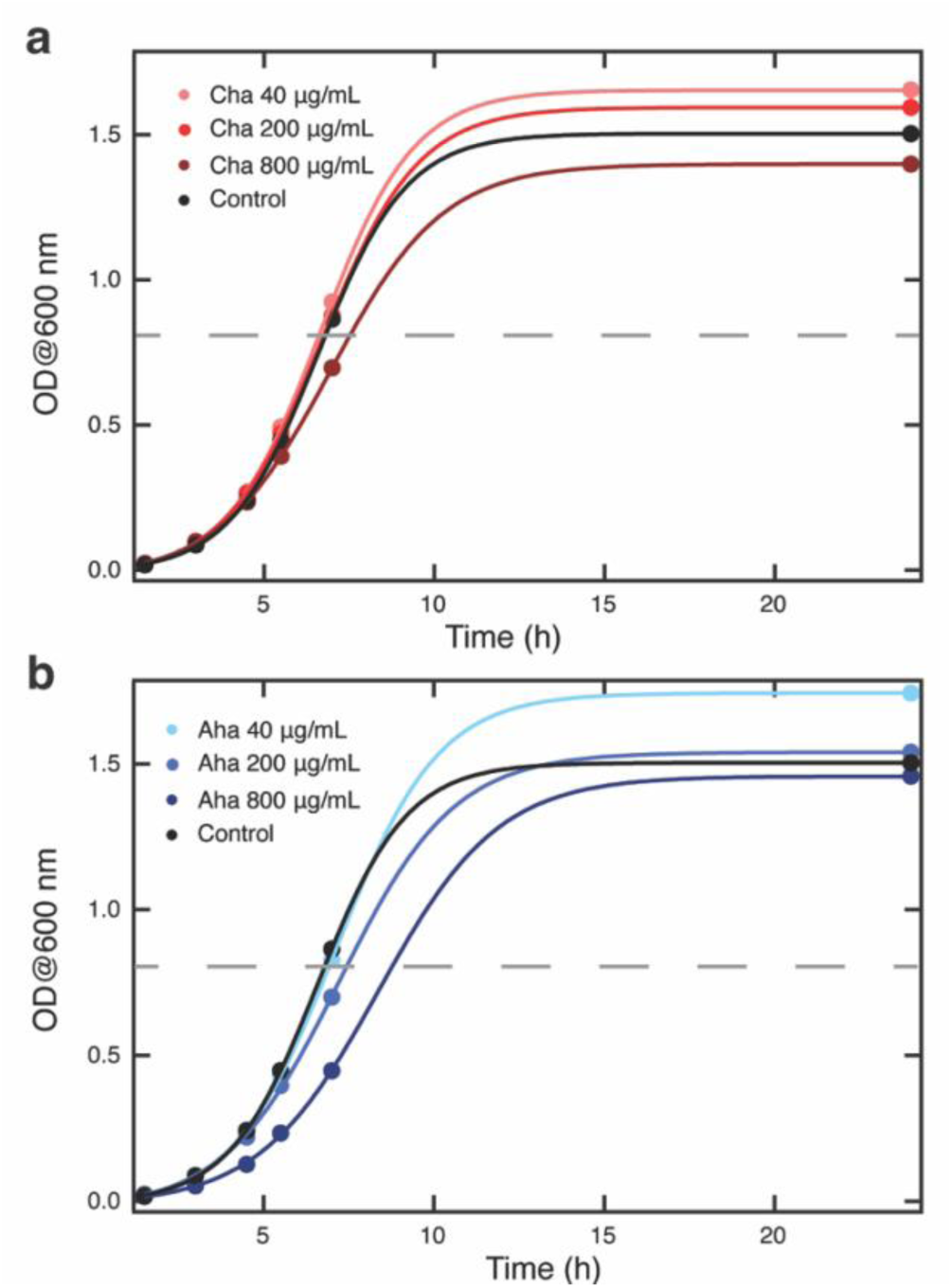
Effects of *L-*cyanohomoalanine (Cha) and *L-*azidohomoalanine (Aha) on prototrophic *E. coli* growth. Growth curves from measuring the optical density at 600 nm (OD@600 nm) for BL21(DE3) *E. coli* grown in M9 minimal media with (a) Cha or (b) Aha supplemented at 0 (control), 40, 200, or 800 μg/mL. 20 mL cultures were grown in 50 mL falcon tubes at 37 °C with 200 rpm shaking. Plots are fit with a sigmoid. Dashed lines indicate an OD@600 nm of 0.8. Data visualized in IgorPro and figure assembled in Adobe Illustrator.

The incorporation of Met, Aha, and Cha into a modified super folder green fluorescent protein (sfGFP) was evaluated in a Met auxotroph *E. coli* cell line (B834 DE3) that does not natively produce Met using the method depicted in Fig. 1. Briefly, in line with previous work (Fig. 1a),^17^ a single colony was selected from a transformation and grown overnight in LB media. The overnight culture was transferred to a large culture of M9 minimal media supplemented with Met and allowed to grow until reaching an OD600 of 0.8-1. The cells were pelleted and starved for 30 minutes before protein expression was induced in fresh M9 minimal media supplemented with Met, Aha, or Cha (Fig. 1b). The sfGFP mutant contains only two Met residues, an N-terminal methionine to initiate protein expression and a single other methionine at either position 150 or 234 (Fig. 1c-d) so that the site-specific effects of the ncAA could be evaluated. The second residue of the protein was mutated to alanine to promote cleavage of the N-terminal residue (Fig. 1c).^30^

To determine whether Met, Aha, and Cha were successfully incorporated into sfGFP, the *E. coli* growths were assessed using expression gels (Fig. 3) and fluorometry (Fig. S2). As the *E. coli* used are Met auxotrophs, protein should not be produced in the absence of Met or a Met analog.^32^ However, as the cells are initially cultured in Met containing media, trace Met may remain. Additionally, the *E. coli* may scavenge Met from recycled proteins produced prior to induction. As such, levels of sfGFP expression were compared to both pre-induction (i.e., leaky expression) and 16 hr post-induction cultures not supplemented with Met, Aha, or Cha (Fig. 3 lanes 1-4). A representative gel is shown in Fig. 3a and Fig. 3b shows the average quantification across four replicates. No sfGFP is detected in the pre-induction lanes (Fig. 3 lanes 1-2). However, as suspected, sfGFP is observed post-induction in cells not supplemented with Met, Aha, or Cha (Fig. 3 lanes 3-4), arising either from residual Met from pre-induction or recycling Met from other proteins. Compared to expression of sfGFP in cells fed Met, Aha labeled sfGFP appears to express at ≈2/3^rd^ the level while Cha labeled sfGFP expressed at ≈1/3^rd^ the level. Cha expression levels are slightly above those of control cells not supplemented with Met, Aha, or Cha (Fig. 3 lanes 3-4 vs lanes 9-10), suggesting low but present levels of Cha incorporation. The expression levels of sfGFP in *E. coli* growths were also assessed using sfGFP as a proxy for expression levels. Because sfGFP is a fluorescent protein its fluorescence can be uniquely detected within the *E. coli* cytosol. Fluorometry of *E. coli* growths (Fig. S2) produced similar results to the expression gels (Fig. 3) for both 150 and 234 sfGFP, with the fluorescence intensity of cells fed Met > Aha > Cha > control cells not supplemented post-induction. In all of these measurements, Cha sfGFP fluorescence is slightly higher than the control (Fig. S2 red, green). Taken together, our expression gels and fluorometry suggest that Cha is recognized by wild-type methionyl-tRNA synthetase.

**Figure 3.**
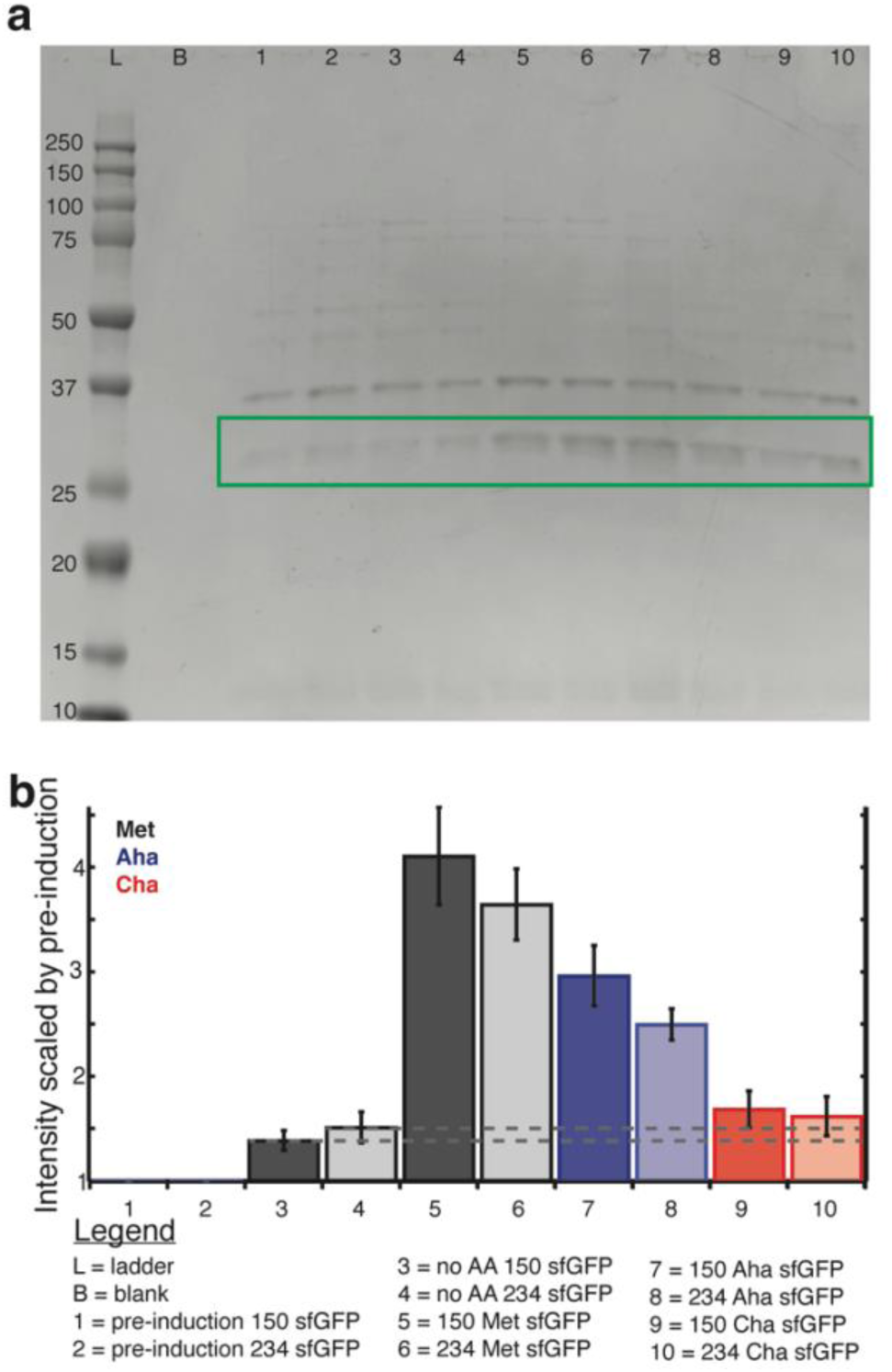
Expression of superfolder GFP (sfGFP) mutants containing *L-*methionine (Met), *L-*cyanohomoalanine (Cha), or *L-*azidohomoalanine (Aha). (a) SDS-PAGE gel of lysates of *E. coli* expressions of sfGFP mutants. From left to right the lanes are L. Precision Plus Protein Dual Color Standard, B. blank, 1. pre-induction of 150 sfGFP, 2. pre-induction of 234 sfGFP, 3. 16 hr post-induction of 150 sfGFP with no amino acids supplemented, 4. 16 hr post-induction of 234 sfGFP with no amino acids supplemented, 5. 16 hr post-induction of 150 GFP with Met supplemented, 6. 16 hr post-induction of 234 sfGFP with Met supplemented, 7. 16 hr post-induction of 150 sfGFP with Aha supplemented, 8. 16 hr post-induction of 234 sfGFP with Aha supplemented, 9. 16 hr post-induction of 150 sfGFP with Cha supplemented, and 10. 16 hr post-induction of 234 sfGFP with Cha supplemented. Samples were diluted to identical OD values before lysis. The green box highlights the ≈27 kDa band assigned to sfGFP. (b) Average intensity of the sfGFP bands of four replicate gels, normalized by the protein band at 36 kDa, and then normalized again by the respective pre-induction bands (lanes 1 and 2). Columns 1-10 correspond to the lanes labeled 1-10 on the gel (a). Error bars represent standard error. Lanes are colored by methionine analog: Met (black), Aha (red), and Cha (blue), and shaded according to sfGFP variant: 150 (dark), 234 (light). Dashed lines included to guide the eye. Gels were analyzed in Fiji^31^ and data was visualized in IgorPro by averaging together the results from four replicate gels. Figure assembled in Adobe Illustrator.

To more specifically confirm incorporation of the ncAAs into sfGFP, the proteins were purified, digested with trypsin, and subjected to liquid chromatography-mass spectrometry (LC-MS/MS). Incorporation of Cha and Aha were confirmed in both 150 and 234 sfGFP variants via identification of relevant tryptic peptides (Fig. S3). Labeling efficiency was determined in duplicates by summing the area under the curve of the extracted ion chromatogram (XIC) traces of each labeled tryptic fragment (allowing for two missed cleavages) with +2, +3 and +4 charge and ^12^C, ^13^C and double ^13^C (an example XIC is shown in Fig. S4). To confirm mass shifts did not come from an artefact we performed this analysis on control 150 Met sfGFP and 234 Met sfGFP. Labeling efficiencies are summarized in Table S1. This data should not be taken as fully quantitative as it does not account for differences in ionizability between Aha, Cha, and Met labeled peptides. For 234 Cha sfGFP, Cha labeling occurred at 10% and 4% of positions for trial 1 and trial 2, respectively. For 234 Aha sfGFP, Aha labeling occurred at 49% and 68%. In control 234 Met sfGFP and 150 Met sfGFP, Aha and Cha labeling (i.e. artefacts) was ≤1.5%. In 150 Aha sfGFP, Aha labeling efficiency was similar to 234 Aha sfGFP with 76% and 50% labeled in trial 1 and 2, respectively. However, 234 Cha sfGFP had Cha labeling in only 1.9% and 1.2% of positions, near the artefact level. Surprisingly, 3% and 8% Aha were identified in 150 Cha sfGFP trial 1 and 2, respectively. These cultures were not fed Aha and cannot manufacture Aha on their own. Further inspection revealed that the mass difference of oxidized Cha (oxCha) compared to Met (−4.99349 Da) is nearly identical to the mass difference of Aha compared to Met (−4.98634 Da). Thus, it is possible that the solvent exposed 150 position of Cha is modified and Cha labeling efficiency in 150 Cha sfGFP is closer to the sum of oxCha and Cha, i.e. 5-9%, which corresponds with the 234 Cha sfGFP Cha labeling rate.

We further verified incorporation of Cha and Aha in sfGFP via tandem mass spectrometry (MS2) by inspecting the fragments and confirming the b+ and/or the y+ fragment ions of precursor peptides contained the substituted amino acid. For 234 sfGFP (Fig. 4), y+ fragment ions y7+, y8+, y9+, y10+, y11+, y12+, y13+, and y14+ have a shift in mass consistent with incorporation of the ncAA at the y6 position of the precursor peptide fragment (Table S2-3). However, none of the b+ fragments identified in Fig. 4 are long enough to confirm ncAA incorporation at the b18 position. Incorporation of Aha and Cha was also confirmed with MS2 fragmentation in 150 sfGFP (Fig. S5a-b, Table S4-5). However, the intensity of 150 Cha XIC and MS2 fragments was low, likely due to the oxidation of Cha (Table S1, Figure S3). Indeed, when instead inspecting for oxCha at position 150, incorporation of oxCha can be confirmed in the fragmentation patterns (Fig. S4c, Table S6). Additionally, LC-MS/MS confirmed cleavage of the N-terminal methionine as no fragments containing the N-terminal methionine were visualized (Table S7). Thus Cha, like Aha,^30,33^ is likely a substrate of methionine aminopeptidase (MAP) as no Cha N-terminal residue was detected. This is important for site specificity in downstream applications of Cha for bioorthogonal chemistry or vibrational and MS probes.

**Figure 4.**
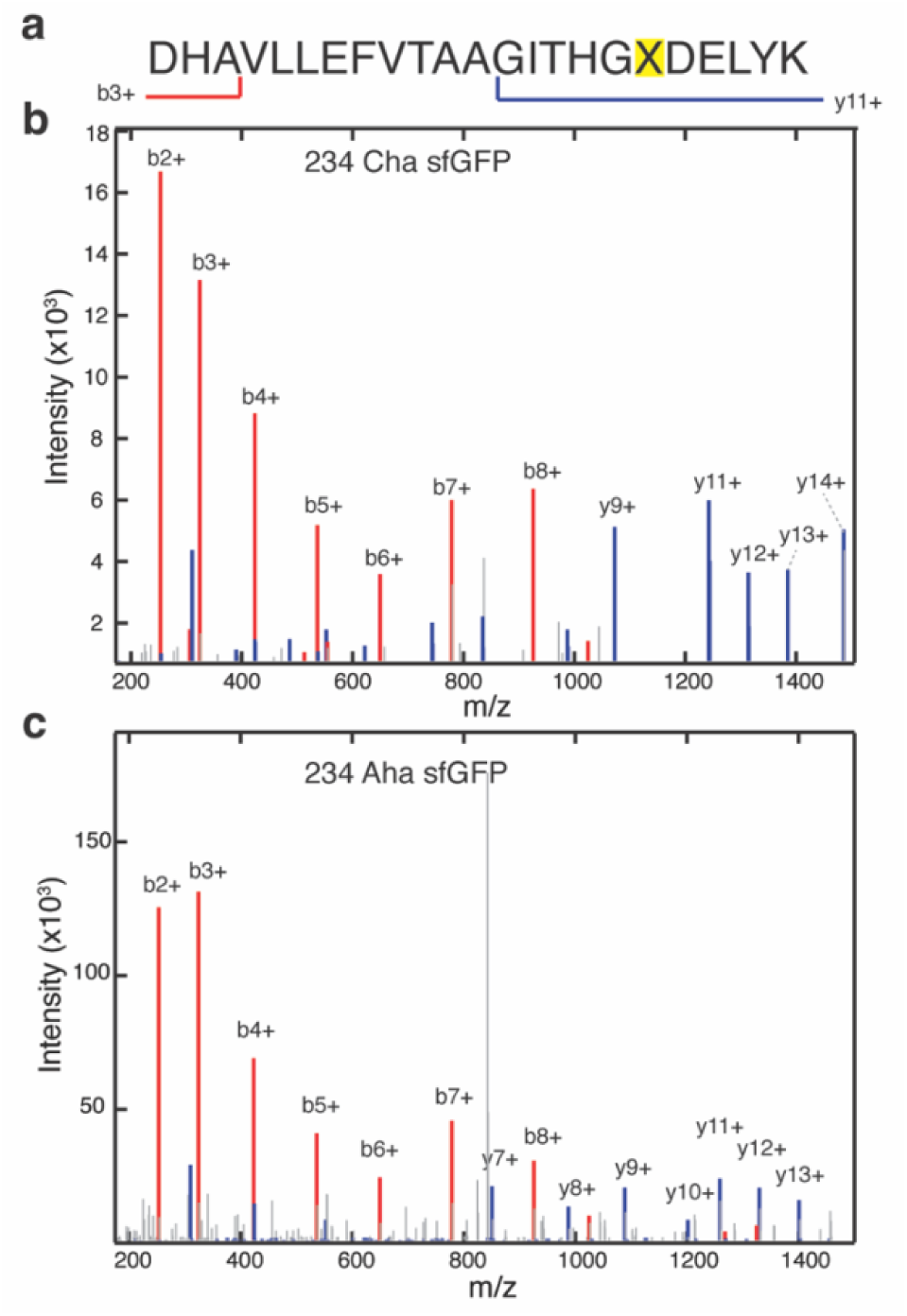
Tandem mass spectrometry (MS2) incorporation of *L-*cyanohomoalanine (Cha) and *L-*azidohomoalanine (Aha) into superfolder GFP (sfGFP). (a) Example precursor peptide sequence of sfGFP containing the 234 ncAA incorporation site, highlighted X, selected for fragmentation. MS2 fragmentation spectra of the (a) precursor peptide for (b) 234 Cha sfGFP and (c) 234 Aha sfGFP. b+ ions denote N-terminal fragments (red) and y+ ions denote C-terminal fragments (blue). Corresponding lists of masses and sequences can be found in Tables S2-3 with identified fragments bolded. Data visualized in IgorPro and figure assembled in Adobe Illustrator.

Having confirmed the incorporation of Cha and Aha to sfGFP, we investigated whether the ncAAs disrupted the stability of sfGFP via temperature melts (Fig. S6). The average melting temperatures for the sfGFPs are reported in Table 1. No change in melting temperature was observed for any of the 150 sfGFP proteins (Fig. 5a, Table 1, Table S8). This is anticipated as position 150 is solvent exposed, with the sidechain facing out of the β-barrel (Fig. 1d), substitution with Cha and Aha should not affect protein stability. Incorporation of Cha to 234 sfGFP resulted in a destabilization with borderline statistical significance (p=0.053 by one-tailed student t-test) while incorporation of Aha had no effect on 234 sfGFP stability (Fig. 5b, Table S8). Destabilization of Cha at position 234 is surprising because position 234 is in the disordered C-terminal tail of sfGFP (Fig. 1d). One possibility is that packing of the C-terminal tail of sfGFP against the β-barrel destabilizes it. As nitrile moieties are more polar than azide moieties, burying the nitrile would have a higher thermodynamic cost which could result in a minor disruption. Thus, it is important to check the effect of ncAA on the protein properties of interest, even if the probe is small and is incorporated far from the protein core.

**Figure 5.**
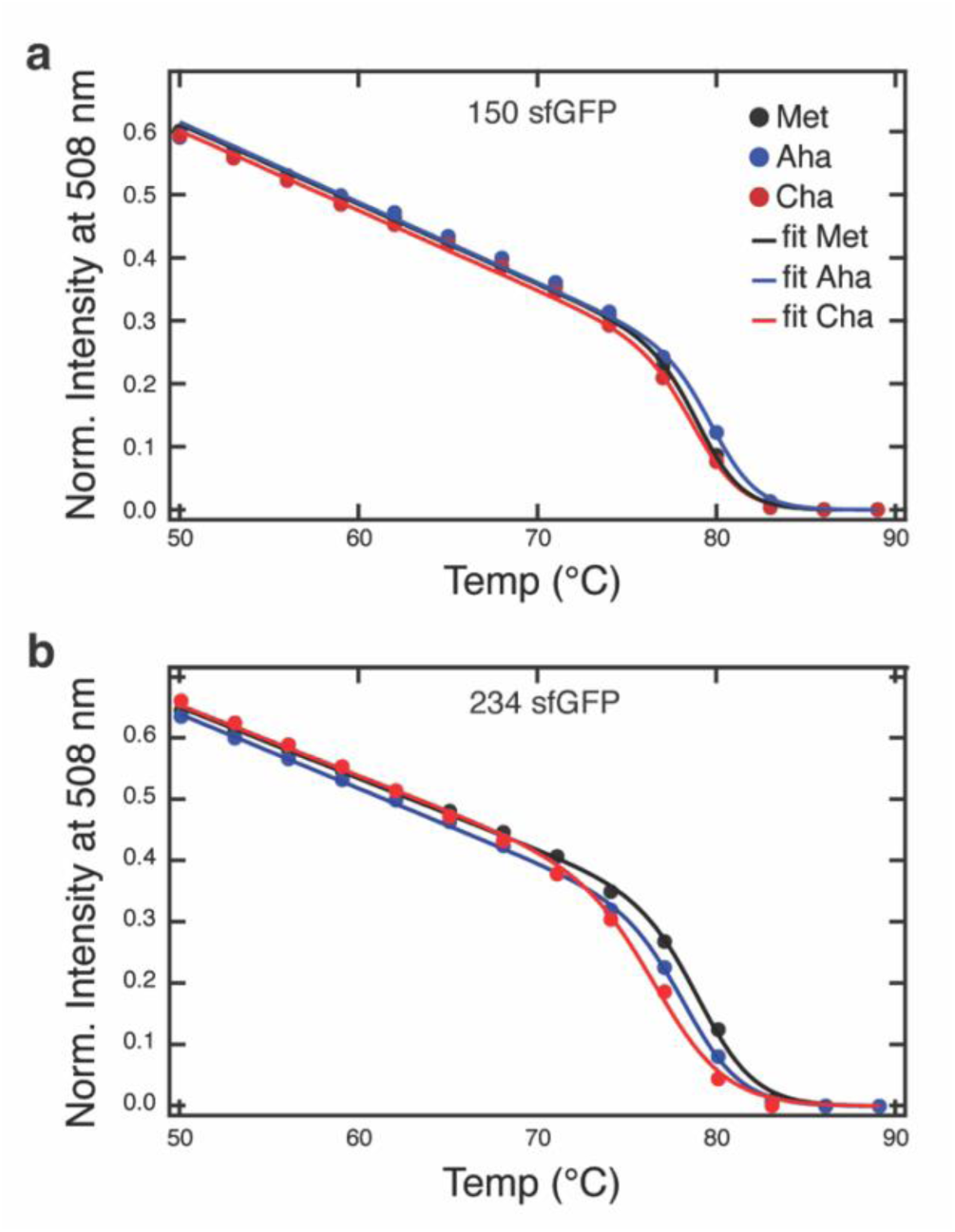
Representative melting curves of sfGFP variants monitored from 20 to 89 °C in 3 °C steps. Single, representative plots of the change in intensity at 508 nm with increasing temperature for (a) 150 sfGFP variants and (b) 234 sfGFP variants. [protein] = 5 μΜ in 10 mM sodium phosphate buffer, pH 7.0. Met is colored black, Aha is colored blue, and Cha is colored red. Plots are fit with a two-state denaturation (Eq. S1) overlaid on the data. Data visualized in Igor Pro and figure assembled in Adobe Illustrator.

**Table 1.** Average melting temperatures of sfGFP variants.

| **Protein** | **T<sub>m</sub> (°C)** |
| --- | --- |
| **150 Met sfGFP** | 80 ± 1 |
| **150 Aha sfGFP** | 81 ± 1 |
| **150 Cha sfGFP** | 79 ± 2 |
| **234 Met sfGFP** | 79.4 ± 0.9 |
| **234 Aha sfGFP** | 78 ± 2 |
| **234 Cha sfGFP** | 76 ± 3 |

## Conclusion

We demonstrate incorporation of a novel ncAA, Cha, into sfGFP, and compare it against a known ncAA, Aha. Importantly, Cha incorporation does not destabilize sfGFP significantly at either position tested, though possible disruption in hydrophobic regions of proteins should be considered when using the ncAA (or any ncAA) for downstream applications. Competition from canonical Met was minimized by performing experiments in Met auxotroph *E. coli.* We find that Cha serves as a substrate for metRS and successfully incorporates, although expression gels, fluorometry, and LC-MS/MS data suggest that Cha incorporation is less efficient than for Aha. However, Cha is less disruptive to *E. coli* growth than Aha, suggesting that low incorporation is not due to cytotoxicity. Due to its small size and similarity to Hpg, it is also unlikely that Cha has a structural incompatibility with translational machinery that causes steric hinderance and ribosomal stalling.^34^ Thus, the lower incorporation of Cha to sfGFP is likely due to low recognition by the WT methionyl-tRNA synthetase compared to Met or Aha. This is surprising given the similarity to Cha to Hpg and Aha.

In the future, Cha may be useful for bioorthogonal chemical reactions that make use of the nitrile moiety, especially in cases where multiple click probes are desirable.^24^ The reactions of Aha with an alkyne and Cha with an allene are orthogonal. Cha also has potential as a MS probe orthogonal to Aha and Hpg due to their distinct masses, allowing for pulse-chase experiments with three separate pulse events, tracking protein expression over time.^6^ Importantly, Cha oxidation should be considered during mass spectrometry experiments, especially when used in tandem with Aha as it can lead to very similar masses. Nitrile moieties are also suitable probes for both Raman and IR spectroscopy and imaging as they are highly sensitive to local environement.^19,20^ However, the current levels of Cha incorporation are likely too low for it to be detected as a vibrational probe. Future work evolving *E. coli* could lead to increased incorporation for this bioorthogonal vibrational probe. Further, isotopic labeling of Cha could create many vibrational and MS ‘colors’ for the probe.^25^ Taken together, Cha is a non-toxic ncAA probe with potential as both a vibrational and mass spectrometry label and a chemical handle.

## ASSOCIATED CONTENT

**Supporting Information**. Cha SI

Materials and methods, fluorescence spectra of *E. coli* growths of the sfGFP mutants (Fig. S1), MS of mutant sfGFp variants (Fig. S2), example XIC (Fig. S3), quantification of Aha and Cha labeling (Table S1), expected fragmentation for representative peptides from 234 Cha and Aha sfGFP (Table S2-3), MS2 of 150 Aha, 150 Cha and 150 oxidized Cha (Fig. S4), expected fragmentation for representative peptides from 150 Aha and Cha, and oxidized Cha sfGFP (Table S4-6), full spectra of 234 Cha sfGFP thermal melt (Fig. S5), melting temperatures of each sfGFP variant (Table S8), and sfGFP variant genes (Table S9).

N-terminal fragments

List of identified N-terminal peptides (Table S7)

## Author Contributions

The manuscript was written through contributions of all authors. All authors have given approval to the final version of the manuscript.

## Funding Sources

This work was supported by a Beckman Young Investigator Award from the Arnold and Mabel Beckman Foundation (https://dx.doi.org/10.13039/100000997). S.O.S. was partially supported by the NIH under training grant T32 GM008283 and a National Science Foundation Graduate Research Fellowship under grant DGE2139841. The authors also thank the MS & Proteomics Resource at Yale University for providing the necessary mass spectrometers and the accompanying biotechnology tools, funded in part by the Yale School of Medicine and by the Office of the Director, National Institutes of Health (S10OD02365101A1, S10OD019967, and S10OD018034). The funders had no role in study design, data collection and analysis, decision to publish, or preparation of the manuscript.

## ACKNOWLEDGMENT

The authors thank the MS & Proteomics Resource at Yale University for providing the necessary mass spectrometers and the accompanying biotechnology tools.

## ABBREVIATIONS

Aha: L-azidohomoalanine
Cha: L-cyanohomoalanine
CuAAC: copper-catalyzed azide-alkyne cycloaddition
GCE: genetic code expansion
Hpg: L-homopropargylglycine
IR: infrared
IPTG: Isopropyl β-D-1-thiogalactopyranoside
LB: Luria broth
LC-MS: liquid chromatography mass-spectrometry
MAP: methionine aminopeptidase
Met: L-methionine
metRS: methionyl-tRNA synthease
MS: mass spectrometry
MS2: tandem mass spectrometry
ncAA: non-canonical amino acid
OD: optical density
O/N: overnight pMet oxCha oxidized Cha; L-photo-methionine
pCNPhe: p-cyano-L-phenylalanine
SeMet: selenomethionine
sfGFP: superfolder green fluorescent protein
XIC: extracted ion chromatogram

## TOC Figure

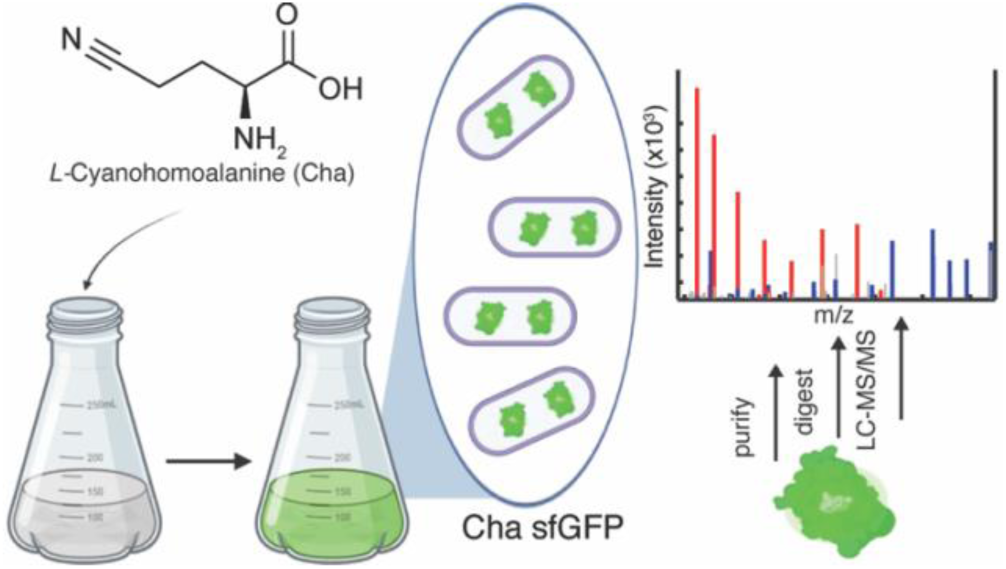

## Notes

### Competing Interest Statement

The authors have declared no competing interest.

## References.

(1) Feng, R.; Wang, M.; Zhang, W.; Gai, F. Unnatural Amino Acids for Biological Spectroscopy and Microscopy. Chem. Rev. 2024, 124 (10), 6501–6542. 10.1021/acs.chemrev.3c00944.

(2) Zhang, W. H.; Otting, G.; Jackson, C. J. Protein Engineering with Unnatural Amino Acids. Curr Opin Struct Biol 2013, 23, 581–587.

(3) Mitra, N. Incorporating Unnatural Amino Acids into Recombinant Proteins in Living Cells. *Mater*. Methods 2023.

(4) Costello, A.; Peterson, A. A.; Chen, P.-H.; Bagirzadeh, R.; Lanster, D. L.; Badran, A. H. Genetic Code Expansion History and Modern Innovations. Chem. Rev. 2024, 124 (21), 11962–12005. 10.1021/acs.chemrev.4c00275.

(5) Liu, C. C.; Schultz, P. G. Adding New Chemistries to the Genetic Code. Annu. Rev. Biochem. 2010, 79 (Volume 79, 2010), 413–444. 10.1146/annurev.biochem.052308.105824.

(6) Dieterich, D. C.; Link, A. J.; Graumann, J.; Tirrell, D. A.; Schuman, E. M. Selective Identification of Newly Synthesized Proteins in Mammalian Cells Using Bioorthogonal Noncanonical Amino Acid Tagging (BONCAT). Proc. Natl. Acad. Sci. 2006, 103 (25), 9482–9487. 10.1073/pnas.0601637103.

(7) van Hest, J. C. M.; Kiick, K. L.; Tirrell, D. A. Efficient Incorporation of Unsaturated Methionine Analogues into Proteins in Vivo. J. Am. Chem. Soc. 2000, 122 (7), 1282–1288. 10.1021/ja992749j.

(8) Cowie, D. B.; Cohen, G. N. Biosynthesis by *Escherichia Coli* of Active Altered Proteins Containing Selenium Instead of Sulfur. Biochim. Biophys. Acta 1957, 26 (2), 252–261. 10.1016/0006-3002(57)90003-3.

(9) Jecmen, T.; Tuzhilkin, R.; Sulc, M. Photo-Methionine, Azidohomoalanine and Homopropargylglycine Are Incorporated into Newly Synthesized Proteins at Different Rates and Differentially Affect the Growth and Protein Expression Levels of Auxotrophic and Prototrophic E. Coli in Minimal Medium. Int. J. Mol. Sci. 2023, 24 (14), 11779. 10.3390/ijms241411779.

(10) Leahy, D. J.; Hendrickson, W. A.; Aukhil, I.; Erickson, H. P. Structure of a Fibronectin Type III Domain from Tenascin Phased by MAD Analysis of the Selenomethionyl Protein. Science 1992, 258 (5084), 987–991. 10.1126/science.1279805.

(11) Watson, M. D.; Lee, J. C. Genetically Encoded Aryl Alkyne for Raman Spectral Imaging of Intracellular α-Synuclein Fibrils. J. Mol. Biol. 2022, 167716. 10.1016/j.jmb.2022.167716.

(12) Taskent-Sezgin, H.; Chung, J.; Banerjee, P. S.; Nagarajan, S.; Dyer, R. B.; Carrico, I.; Raleigh, D. P. Azidohomoalanine: A Conformationally Sensitive IR Probe of Protein Folding, Protein Structure, and Electrostatics. Angew. Chem. Int. Ed. 2010, 49 (41), 7473–7475. 10.1002/anie.201003325.

(13) Korona, W.; Orzechowska, B.; Si¹ka³a, K.; Nowakowska, A. M.; Pieczara, A.; Buda, S.; Pawlowski, R.; Mlynarski, J.; Baranska, M. Bioorthogonal Raman and IR Probes for Live Cell Metabolomics: A Library. Sens. Actuators B Chem. 2025, 430, 137363. 10.1016/j.snb.2025.137363.

(14) Kirschner, F.; Arnold-Schild, D.; Leps, C.; Łącki, M. K.; Klein, M.; Chen, Y.; Ludt, A.; Marini, F.; Kücük, C.; Stein, L.; Distler, U.; Sielaff, M.; Michna, T.; Riegel, K.; Rajalingam, K.; Bopp, T.; Tenzer, S.; Schild, H. Modulation of Cellular Transcriptome and Proteome Composition by Azidohomoalanine—Implications on Click Chemistry–Based Secretome Analysis. J. Mol. Med. 2023, 101 (7), 855–867. 10.1007/s00109-023-02333-4.

(15) Tivendale, N. D.; Fenske, R.; Duncan, O.; Millar, A. H. In Vivo Homopropargylglycine Incorporation Enables Sampling, Isolation and Characterization of Nascent Proteins from Arabidopsis Thaliana. Plant J. 2021, 107 (4), 1260–1276. 10.1111/tpj.15376.

(16) Landor, L. A. I.; Bratbak, G.; Larsen, A.; Tjendra, J.; Våge, S. Differential Toxicity of Bioorthogonal Non-Canonical Amino Acids (BONCAT) in *Escherichia Coli*. J. Microbiol. Methods 2023, 206, 106679. 10.1016/j.mimet.2023.106679.

(17) Flynn, J. D.; Gimmen, M. Y.; Dean, D. N.; Lacy, S. M.; Lee, J. C. Terminal Alkynes as Raman Probes of α-Synuclein in Solution and in Cells. *Chembiochem Eur*. J. Chem. Biol. 2020. 10.1002/cbic.202000026.

(18) Sagawa, N.; Shikata, T. Are All Polar Molecules Hydrophilic? Hydration Numbers of Nitro Compounds and Nitriles in Aqueous Solution. Phys. Chem. Chem. Phys. 2014, 16 (26), 13262–13270. 10.1039/C4CP01280A.

(19) Getahun, Z.; Huang, C.-Y.; Wang, T.; De León, B.; DeGrado, W. F.; Gai, F. Using Nitrile-Derivatized Amino Acids as Infrared Probes of Local Environment. J. Am. Chem. Soc. 2003, 125 (2), 405–411. 10.1021/ja0285262.

(20) Andrews, S. S.; Boxer, S. G. Vibrational Stark Effects of Nitriles II. Physical Origins of Stark Effects from Experiment and Perturbation Models. J. Phys. Chem. A 2002, 106 (3), 469–477. 10.1021/jp011724f.

(21) Weaver, J. B.; Kozuch, J.; Kirsh, J. M.; Boxer, S. G. Nitrile Infrared Intensities Characterize Electric Fields and Hydrogen Bonding in Protic, Aprotic, and Protein Environments. J. Am. Chem. Soc. 2022, 144 (17), 7562–7567. 10.1021/jacs.2c00675.

(22) Beauchard, A.; Twum, E. A.; Lloyd, M. D.; Threadgill, M. D. *S*-2-Amino-4-Cyanobutanoic Acid (β-Cyanomethyl-l-Ala) as an Atom-Efficient Solubilising Synthon for l-Glutamine. Tetrahedron Lett. 2011, 52 (41), 5311–5314. 10.1016/j.tetlet.2011.08.017.

(23) Marzi, M.; Vakil, M. K.; Bahmanyar, M.; Zarenezhad, E. Paxlovid: Mechanism of Action, Synthesis, and In Silico Study. BioMed Res. Int. 2022, 2022, 7341493. 10.1155/2022/7341493.

(24) Paioti, P. H. S.; Lounsbury, K. E.; Romiti, F.; Formica, M.; Bauer, V.; Zandonella, C.; Hackey, M. E.; del Pozo, J.; Hoveyda, A. H. Click Processes Orthogonal to CuAAC and SuFEx Forge Selectively Modifiable Fluorescent Linkers. Nat. Chem. 2024, 16 (3), 426–436. 10.1038/s41557-023-01386-9.

(25) Bazewicz, C. G.; Lipkin, J. S.; Smith, E. E.; Liskov, M. T.; Brewer, S. H. Expanding the Utility of 4-Cyano-l-Phenylalanine As a Vibrational Reporter of Protein Environments. J. Phys. Chem. B 2012, 116 (35), 10824–10831. 10.1021/jp306886s.

(26) Davis, C. BioRender. Biorender. https://app.biorender.com/citation/6a0b70e588db31d12dd27c26 (accessed 2026-05-18).

(27) Abramson, J.; Adler, J.; Dunger, J.; Evans, R.; Green, T.; Pritzel, A.; Ronneberger, O.; Willmore, L.; Ballard, A. J.; Bambrick, J. et. al. Accurate Structure Prediction of Biomolecular Interactions with AlphaFold 3. Nature 2024, 630 (8016), 493–500. 10.1038/s41586-024-07487-w.

(28) Brysk, M. M.; Ressler, C. γ-Cyano-α-l-Aminobutyric Acid: A New Product Of Cyanide Fixation in *Chromobacterium Violaceum*. J. Biol. Chem. 1970, 245 (5), 1156–1160. 10.1016/S0021-9258(18)63301-0.

(29) Rothenberg, D. A.; Taliaferro, J. M.; Huber, S. M.; Begley, T. J.; Dedon, P. C.; White, F. M. A Proteomics Approach to Profiling the Temporal Translational Response to Stress and Growth. iScience 2018, 9, 367–381. 10.1016/j.isci.2018.11.004.

(30) Wingfield, P. N-Terminal Methionine Processing. Curr. Protoc. Protein Sci. 2017, 88, 6.14.1–6.14.3. 10.1002/cpps.29.

(31) Schindelin, J.; Arganda-Carreras, I.; Frise, E.; Kaynig, V.; Longair, M.; Pietzsch, T.; Preibisch, S.; Rueden, C.; Saalfeld, S.; Schmid, B. et. al. Fiji: An Open-Source Platform for Biological-Image Analysis. Nat. Methods 2012, 9 (7), 676–682. 10.1038/nmeth.2019.

(32) Wood, W. B. Host Specificity of DNA Produced by *Escherichia Coli*: Bacterial Mutations Affecting the Restriction and Modification of DNA. J. Mol. Biol. 1966, 16 (1), 118-IN3. 10.1016/S0022-2836(66)80267-X.

(33) Wang, A.; Winblade Nairn, N.; Johnson, R. S.; Tirrell, D. A.; Grabstein, K. Processing of N-Terminal Unnatural Amino Acids in Recombinant Human Interferon-β in Escherichia Coli. ChemBioChem 2008, 9 (2), 324–330. 10.1002/cbic.200700379.

(34) Saleh, A. M.; Wilding, K. M.; Calve, S.; Bundy, B. C.; Kinzer-Ursem, T. L. Non-Canonical Amino Acid Labeling in Proteomics and Biotechnology. J. Biol. Eng. 2019, 13, 43. 10.1186/s13036-019-0166-3.

