## Supplementary material for "Expanding the methionine toolkit: *L*-cyanohomoalanine as a multifunctional analog": Methods and SI Figures/Tables

**Affiliation**
Department of Chemistry, Yale University, New Haven, Connecticut 06511, United States

**This file includes:**

Materials and Methods

Supplementary Fig. 1 to Supplementary Fig. 5

Tables S1 to S6

Tables S8 and S9

**Other Supplementary Materials for this manuscript include the following:**

Table S7

**Materials.** Unless otherwise stated, all materials and chemicals were sourced from Sigma-Aldrich.

***E. coli* growth.** pUC19 was transformed into BL21(DE3) competent *E. coli* (New England BioLabs, Ipswich, MA) to impart ampicillin resistance. All growths were done under the selective pressure of 100 mg/L ampicillin. One colony was selected from the transformation and used to inoculate 5 mL of LB, grown overnight at 37°C. 200 µL of growth was used to inoculate 20 mL of M9 minimal media and the samples grown with or without Aha or Cha at 40, 200, or 800 mg/L. Samples were grown at 37°C with shaking at 200 rpm and 1 mL removed to measure the optical density at 600 nm (OD600) via UV-Vis spectrometry (Shimadzu, Kyoto, Japan) at the indicated time points. Resulting growth curved were fit in Igor Pro 8 (Wavemetrics, Lake Oswego, OR) using the sigmoid quick fit option.

**Recombinant Protein Expression and Purification.** The sfGFP gene variants (Table S9), containing a C-terminal 6x His tag, were inserted into pDream2.1 vectors (GenScript Biotech, Piscataway, NJ) containing ampicillin resistance. All growths were done under the selective pressure of 100 mg/L ampicillin. The labeled proteins were expressed from methionine auxotroph B834(DE3) competent *E. coli* (Novagen, Madison, WI) following protocols adapted from Flynn et al.[1] Briefly, one colony was selected from a transformation and used to inoculate 5 mL of LB, grown overnight at 37°C. The overnight culture was added to 1 L complete M9 minimal media supplemented with 40 mg/L methionine and the cells grown at 37 °C until the OD600 reached 0.8-1. Cells were pelleted by centrifugation and resuspended in fresh 1 L M9 media. After a starvation period of 30 minutes at 37 °C, expression was induced with 1 mM isopropyl-β-D-thiogalactopyranoside (IPTG, Inalco, San Luis Obispo, CA) and *L*-azidohomoalanine (Aha) or *L*-cyanohomoalanine (Cha, (S)-2-Amino-4-cyanobutanoic acid, Benchchem, Austin, TX) were added to 40 mg/L. Purified control 150 Met and 234 Met sfGFP were grown by transferring the 5 mL of inoculated LB to 1 L of LB and growing at 37 °C until the OD600 reached 0.6 and then induced with 1 mM IPTG. The induced cells of the control and labeled were grown for 16 h at 20 °C. The cells were harvested by centrifugation, suspended in lysis buffer, and then lysed by sonication. The resulting lysate was centrifuged and filtered prior to being passed through a Ni-NTA column. Fractions containing protein were eluted using imidazole and confirmed by running SDS-PAGE gels. The resulting fractions were then dialyzed into 10 mM sodium phosphate, pH 7.0. Concentration was assessed using the UV absorption at 280 nm and $\varepsilon = 18{,}910\ L^{-1}\ mol^{-1}\ cm^{-1}$. Fluorescence absorbance was not preferred as quantum yield was affected by Aha and Cha incorporation. Protein was concentrated with Amicon 10 kDa spin concentrators to at least 30 µM and frozen with liquid nitrogen and stored at -80 °C until needed.

**Purified Protein fluorescence spectra, melts and analysis.** Fluorescence spectra were collected on a Jasco FP-8500 (Easton, MD). The excitation wavelength was 480 nm, and emission was monitored across 475–700 nm. Melts were performed from 20 to 89°C in 3°C increments, with an equilibration period of 3 min at each temperature. Intensity at 508 nm was plotted against temperature and the resulting curves fit with the following two-state denaturation equation:

$$S(T) = \frac{[\alpha_D + \beta_D(T-273.15)] + [\alpha_N + \beta_N(T-273.15)]e^{\frac{-\Delta G}{RT}}}{\left(1 + e^{\frac{-\Delta G}{RT}}\right)} \qquad \text{(S1)}$$

where $N$ and $D$ denote the native and denatured states, $\alpha_N$ and $\alpha_D$ are the signals from each state at 0°C, $\beta_N$ and $\beta_D$ are the slopes of the baselines for each state, and $\Delta G$ is the free energy of folding. $\alpha_D$ and $\beta_D$ were held at 0 as fluorescence is extinguished in the denatured state. Data was fit using Igor Pro 8. Data was collected in triplicate and averages of melting temperatures are presented with error propagated.

**Protein Expression Gels and *E. coli* Fluorescence Measurements.** Protein for expression gels and fluorometry was expressed similarly to above. One colony was selected from a transformation and used to inoculate 5 mL of LB, grown overnight at 37°C. The overnight culture was added to 25 mL complete M9 minimal media supplemented with 40 mg/L methionine and the cells grown at 37 °C until the OD600 reached 0.8-1. At this point the pre-induction aliquots were taken. The Met controls were at this point induced with 1 mM IPTG. The Aha and Cha samples were pelleted by centrifugation and resuspended in fresh 1-L M9 media. After a starvation period of 30 minutes at 37 °C, expression was induced with 1 mM IPTG and Aha or Cha were added to 40 mg/L. Expression processed for 16 h at 20 °C. 1 mL of cells were aliquoted before induction for each variant and 1 mL after expression. The post-expression samples were diluted to OD at 600 nm of 1 and fluorescence spectra collected on a Jasco FP-8500 (Easton, MD). The excitation wavelength was 480 nm, and emission was monitored across 475–600 nm. Undiluted aliquots were lysed via four freeze-thaw cycles and cell debris removed via centrifugation. Five hundred microliters of the resulting supernatants were concentrated using Amicon 10 kDa spin concentrators to 50 µL. This was combined 1:1 with 2x loading dye (Biorad, Hercules, CA) and imaged via SDS-PAGE.

**Expression Gel Data Analysis.** Gel images were analyzed using the Fiji[2] gel analysis program. Data was normalized by dividing each 27 kDa sfGFP band intensity by the intensity of the band at 37 kDa, a natively occurring E. coli protein. Data was then normalized to the pre-induction control for presentation.

**Protein Digestion.** Purified proteins were prepared for mass spectrometry with S-traps and trypsin digestion according with a S-TrapTM micro spin column digestion kit (Protifi, Fairpoint, NY)

**LCMS Data Acquisition and Analysis.** Liquid chromatography–tandem mass spectrometry (LC–MS/MS) analysis was performed using a Vanquish Neo UHPLC system coupled online to an Q Exactive Orbitrap Mass Spectrometers at the Yale Keck Proteomics Center. Peptide samples were separated over a 182 min gradient with solvent B increased in a stepwise manner (0.5 to 5% over 2 min, 5–25% over 138 min, 25–40% over 25 min, 40–90% over 5 min), followed by a 10 min hold at 90% solvent B to ensure column washing and elution of highly hydrophobic species, followed by 90 to 0.5% over 2 min and column equilibrium. This extended gradient facilitated high-resolution separation of complex peptide mixtures prior to ionization. The mass spectrometer was operated in positive ion mode with full MS scans acquired in the Orbitrap at a resolution of 70,000 over a scan range of 300–1700 m/z. Ion accumulation was controlled with a maximum injection time of 45 ms and an absolute automatic gain control (AGC) target of 3e6. These parameters were chosen to balance sensitivity, mass accuracy, and scan speed. Data-dependent acquisition was employed for MS/MS analysis. Precursor ions were selected using an isolation window of 1.7 m/z and subjected to higher-energy collisional dissociation with a normalized collision energy of 28%. Fragment ions were analyzed in the Orbitrap at a resolution of 17,500, with a maximum injection time of 100 ms and AGC target of 1e5. Dynamic exclusion was set to 30 seconds to minimize repeated sequencing of highly abundant precursor ions, thereby increasing proteome coverage by allowing lower-abundance species to be sampled.

N-terminal Met cleavage analysis: Raw data files were processed using Proteome Discoverer (v3.2, Thermo Fisher Scientific) with the Sequest HT search node for peptide identification. Spectra were searched against a *E. coli* and a customized protein database with tryptic specificity, allowing up to two missed cleavages. Peptide candidates were filtered to a minimum length range of 6 amino acids. Precursor mass tolerance was set to 10 ppm; and the fragment mass tolerance was set to 0.02 Da, consistent with high-resolution Orbitrap detection. Carbamidomethylation of cysteine residues was specified as a static modification, while oxidation of methionine, Met to Aha, and Met to Cha were included as a dynamic modification, with a maximum of three dynamic modifications permitted per peptide. Peptide-spectrum matches (PSMs) were evaluated using a target–decoy strategy to control false discoveries. Identification confidence was filtered using a strict false discovery rate (FDR) threshold of 1% at the peptide and protein levels, and with a relaxed threshold of 5%. Following database searching and validation, protein inference and downstream analyses were conducted within Proteome Discoverer to generate high-confidence protein identifications suitable for further quantitative and functional interpretation.

150 and 234 ncAA incorporation analysis: For quantification and data visualization after fragment identification, raw files were input to Skyline and searched against a custom database containing only the relevant sfGFP variant with tryptic specificity, allowing for up to two missed cleavages with a minimum peptide length of 8 and a maximum of 25. Carbamidomethylation of cysteine residues was specified as a static modification, while oxidation of methionine, Met to Aha, Met to Cha, and Met to oxidated Cha were included as dynamic modifications, with a maximum of three dynamic modifications permitted per peptide. For plotting extracted ion chromatograms (XIC), MS1, and MS2 data Thermo Freestyle (Thermo Fisher), Skyline[3] and fisher_py[4] were used.

**References**


(1) Flynn, J. D.; Gimmen, M. Y.; Dean, D. N.; Lacy, S. M.; Lee, J. C. Terminal Alkynes as Raman Probes of α-Synuclein in Solution and in Cells. Chembiochem 2020. https://doi.org/10.1002/cbic.202000026.

(2) Schindelin, J.; Arganda-Carreras, I.; Frise, E.; Kaynig, V.; Longair, M.; Pietzsch, T.; Preibisch, S.; Rueden, C.; Saalfeld, S.; Schmid, B.; Tinevez, J.-Y.; White, D. J.; Hartenstein, V.; Eliceiri, K.; Tomancak, P.; Cardona, A. Fiji: An Open-Source Platform for Biological-Image Analysis. Nature Methods 2012, 9 (7), 676–682. https://doi.org/10.1038/nmeth.2019.

(3) ProteoWizard. pwiz/pwiz_tools/Skyline at master · ProteoWizard/pwiz. GitHub. https://github.com/ProteoWizard/pwiz/tree/master/pwiz_tools/Skyline (accessed 2026-06-10).
(4) Fisher-Py: This Python Module Allows to Extract Data from the RAW-File-Format Produces by Devices from Thermo Fisher Scientific. https://github.com/ethz-institute-of-microbiology/fisher_py (accessed 2026-05-22).

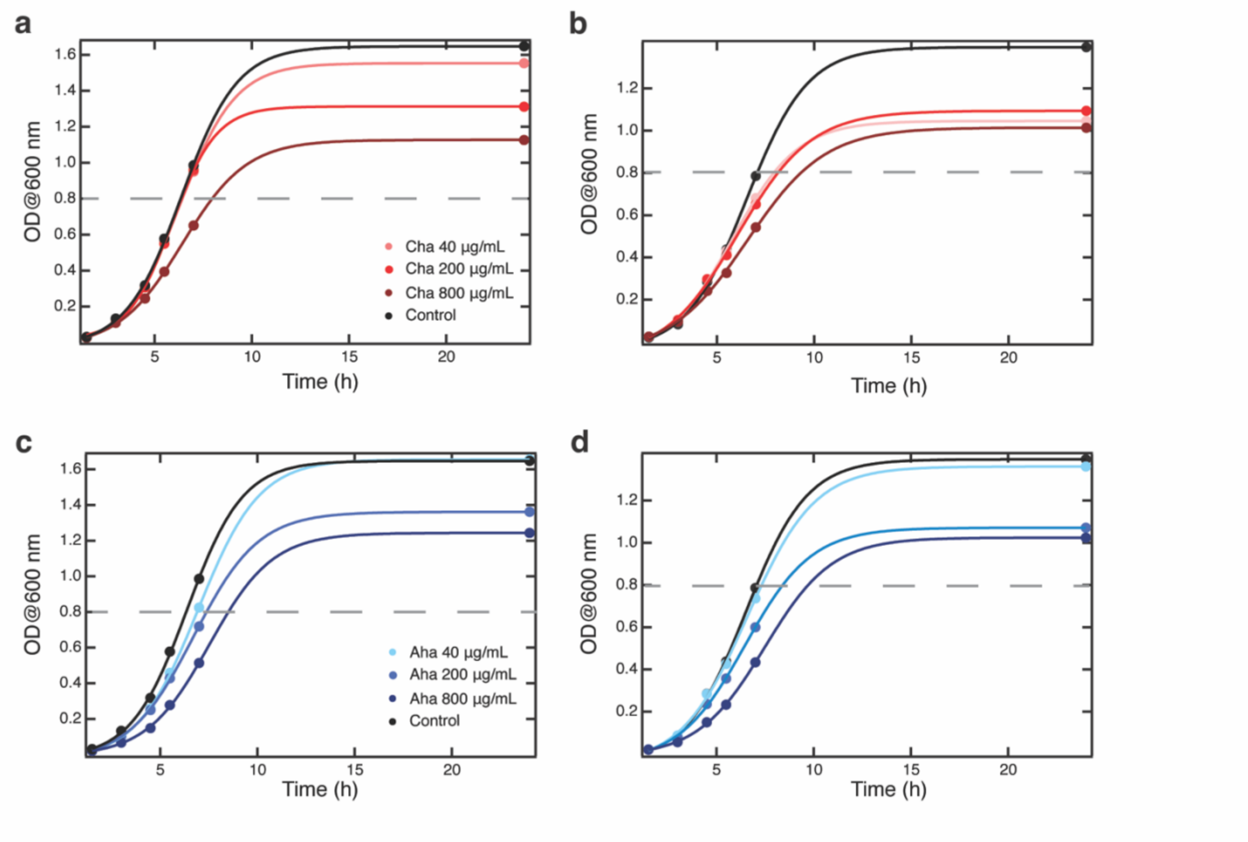


**Figure S1.** Replicate growth curves showing effects of *L*-cyanohomoalanine (Cha) and *L*-azidohomoalanine (Aha) on prototrophic *E. coli* growth. Growth curves from two trials measuring the optical density at 600 nm (OD@600 nm) for BL21(DE3) E. coli grown in M9 minimal media with (a-b) Cha or (c-d) Aha supplemented at 0 (control), 40 , 200, or 800 µg/mL. 20 mL cultures were grown in 50 mL falcon tubes at 37 °C with 200 rpm shaking. Dashed lines indicate an OD@600 nm of 0.8.

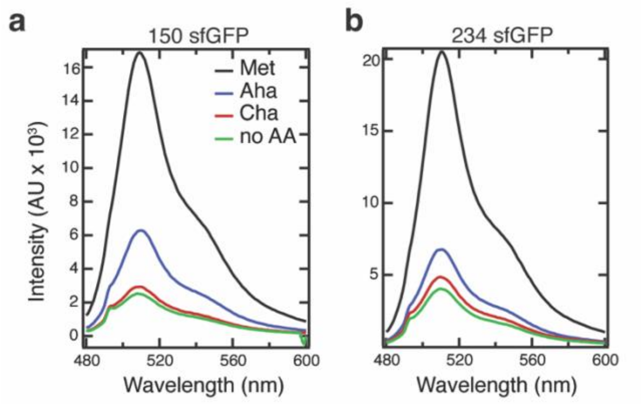


**Figure S2.** Fluorescence spectra of *E. coli* growths of the sfGFP mutants for (a) 150 sfGFP variants and (b) 234 sfGFP variants. The excitation beam was 480 nm. *E. coli* were diluted to OD at 600 nm of 0.5. *E. coli* were fed Met (black), Aha (blue), Cha (red) or nothing (green). Expression was assessed 16 hr post-induction.

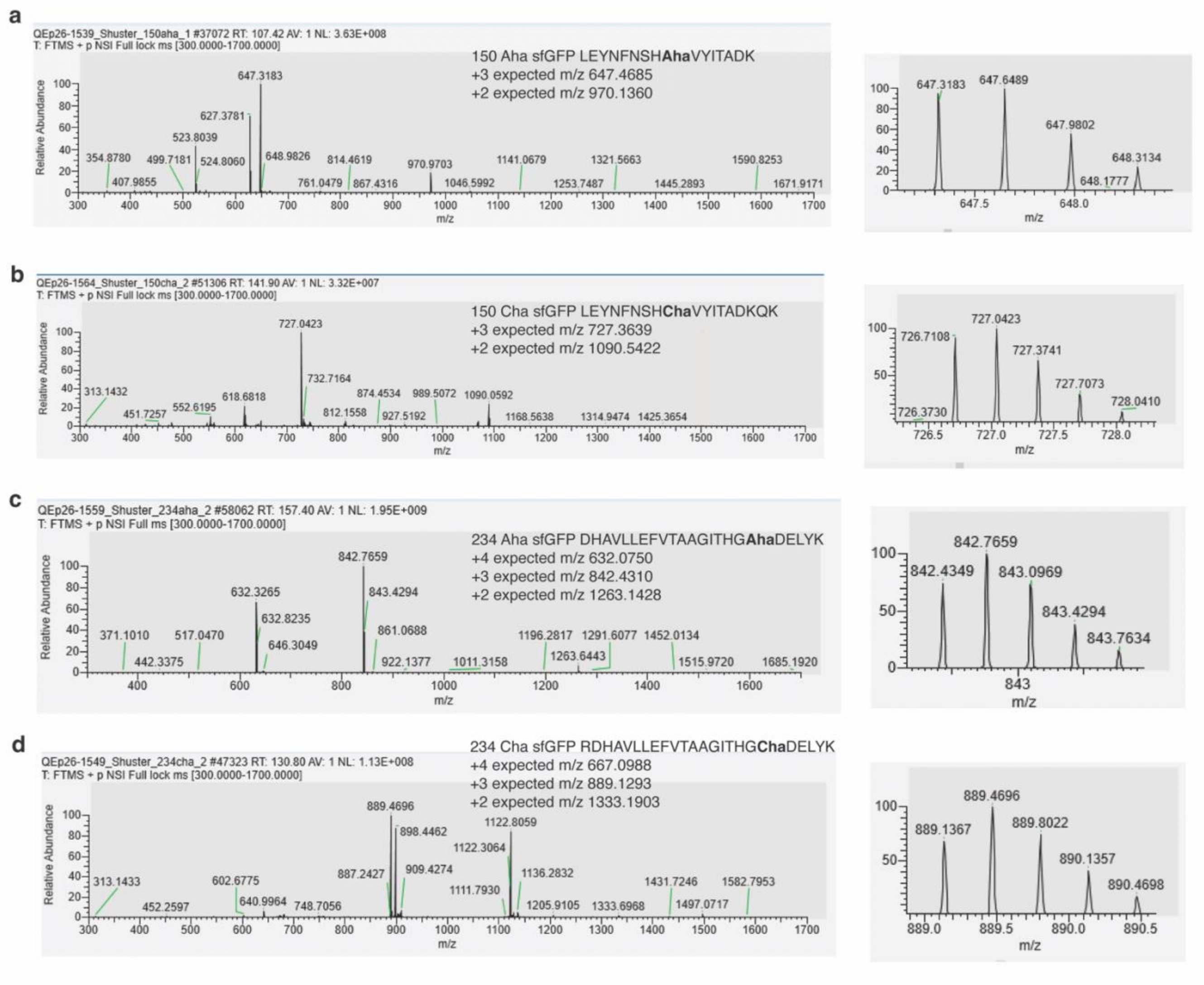


**Figure S3.** Mass Spectrometry (MS1) of mutant sfGFP variants. Representative MS1 of tryptic peptides containing Cha or Aha in (a) 150 Aha, (b) 150 Cha, (c) 234 Aha, and (d) 234 Aha sfGFP. Left shows entire MS range and right provides and enhanced view of one the charge states, showcasing the $^{12}C$, $^{13}C$ and double $^{13}C$ peaks. Data visualized with Thermo Freestyle (Thermo Fisher). Visualized charge states are annotated on the plots. Data is normalized to most intense peak. The runtime in minutes of the MS is noted in the top left of the panels as RT and NL represents the normalization level.

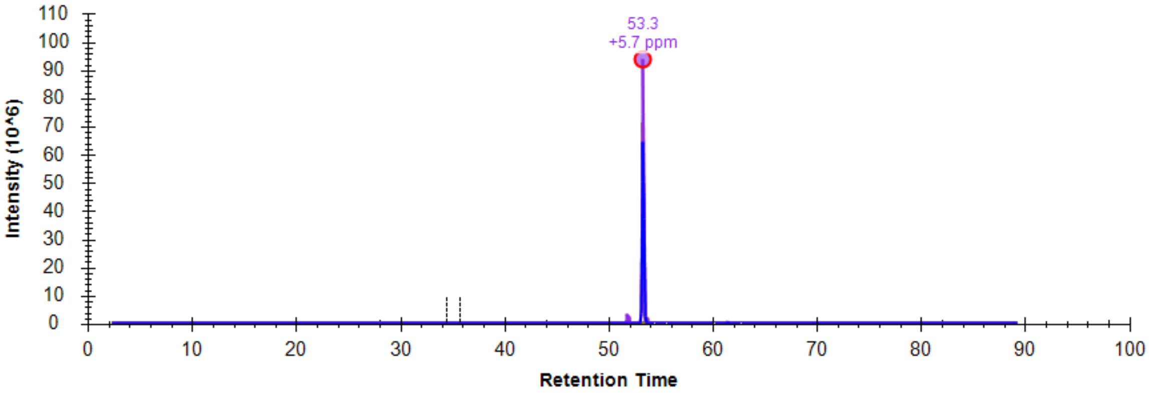


**Figure S4.** Example extracted ion chromatogram (XIC) from 234 Cha sfGFP. The XIC from +3 charge peptide RDHAVLLEFVTAAGITHGChaDELYK at an m/z of 889.1293. Data visualized with Skyline.

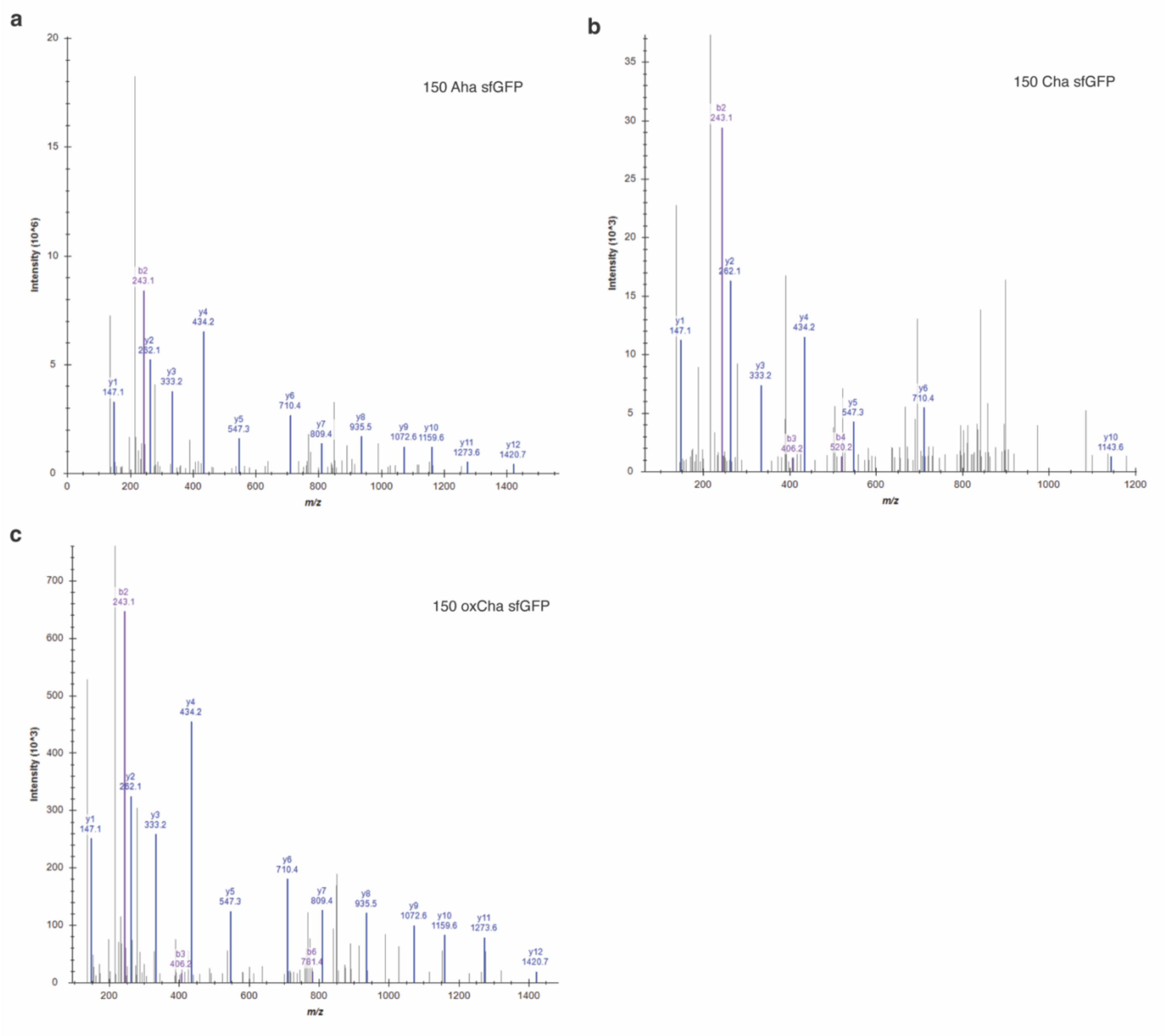


**Figure S5.** Tandem mass spectrometry (MS2) of 150 Aha sfGFP, 150 Cha sfGFP and 150 oxCha sfGFP. MS/MS fragmentation spectra of the precursor peptide for (a) 150 Aha sfGFP, (b) 150 Cha sfGFP, (c) 150 oxidized Cha (oxCha) sfGFP. b+ ions denote N-terminal fragments and y+ ions denote C-terminal fragments (blue). Corresponding lists of masses and sequences can be found in Tables S4-6. Data visualized with Skyline.

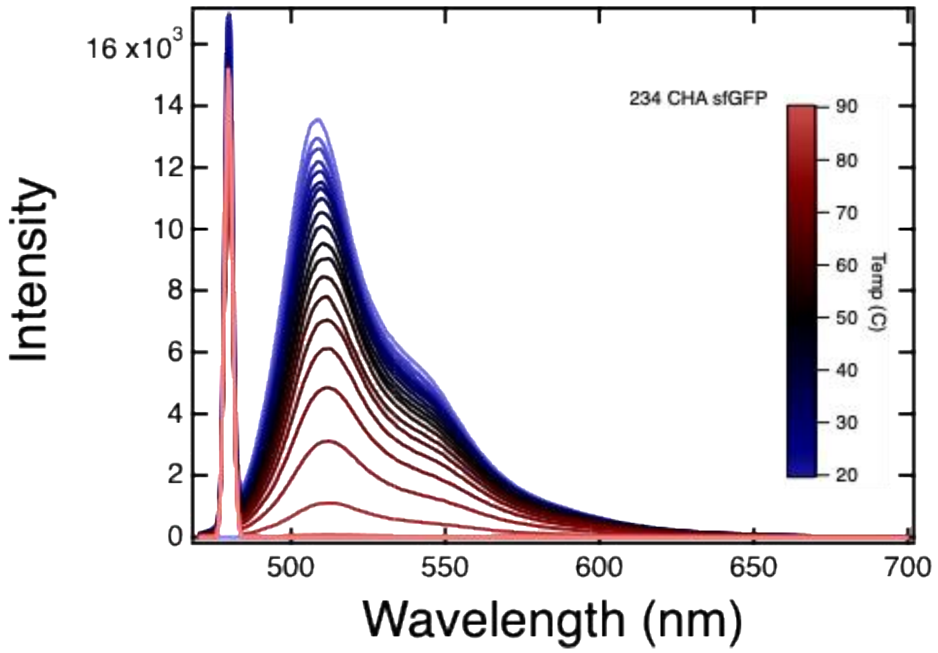


**Figure S6**. Full spectra of 234 Cha sfGFP monitored during a thermal melt. 234 Cha sfGFP (5 μM) in 10 mM sodium phosphate buffer pH 7.0 was monitored from 20 to 89 ºC in 3 ºC steps.

**Table S1.** Extracted Ion Chromatogram (XIC) quantification of Aha and Cha labeling in sfGFP variants

| | % Aha/ oxidized Cha | | % Cha | | % Met | | % oxidized Met | |
|---|---|---|---|---|---|---|---|---|
| | Trial 1 | Trial 2 | Trial 1 | Trial 2 | Trial 1 | Trial 2 | Trial 1 | Trial 2 |
| **150 Met sfGFP** | 0.4 | 1.4 | 0.5 | 1.5 | 90.5 | 91.1 | 8.6 | 6.0 |
| **150 Aha sfGFP** | 76 | 50.4 | 1.0 | 1.4 | 20.9 | 46.1 | 2.0 | 2.1 |
| **150 Cha sfGFP** | 2.9 | 8.1 | 1.9 | 1.2 | 60.7 | 85.5 | 34.5 | 5.2 |
| **234 Met sfGFP** | 0.1 | 0.03 | 1.1 | 1.5 | 90.1 | 84.1 | 8.7 | 14.4 |
| **234 Aha sfGFP** | 49.0 | 67.5 | 0.6 | 1.1 | 35.0 | 28.4 | 15.5 | 3.0 |
| **234 Cha sfGFP** | 0.5 | 0.1 | 9.5 | 3.7 | 44.1 | 79.6 | 45.9 | 16.6 |

**Table S2**. 234 Cha sfGFP mass spec expected fragmentation for example peptide

| | b+ | Seq. | $y^+$ | |
|---|---|---|---|---|
| **1** | 116.03422 | D | | **23** |
| **2** | **253.09313** | H | 2394.24522 | **22** |
| **3** | **324.13025** | A | 2257.18631 | **21** |
| **4** | **423.19866** | V | 2186.1492 | **20** |
| **5** | **536.28272** | L | 2087.08078 | **19** |
| **6** | **649.36679** | L | 1973.99672 | **18** |
| **7** | **778.40938** | E | 1860.91265 | **17** |
| **8** | **925.47779** | F | 1731.87006 | **16** |
| **9** | **1024.54621** | V | 1584.80165 | **15** |
| **10** | 1125.59389 | T | **1485.73323** | **14** |
| **11** | 1196.631 | A | **1384.68555** | **13** |
| **12** | 1267.66811 | A | **1313.64844** | **12** |
| **13** | 1324.68958 | G | **1242.61133** | **11** |
| **14** | 1437.77364 | I | 1185.58986 | **10** |
| **15** | 1538.82132 | T | **1072.5058** | **9** |
| **16** | 1675.88023 | H | **971.45812** | **8** |
| **17** | 1732.9017 | G | **834.39921** | **7** |
| **18** | 1842.94971 | M-Met->Cha | **777.37775** | **6** |
| **19** | 1957.97665 | D | 667.32973 | **5** |
| **20** | 2087.01924 | E | **552.30279** | **4** |
| **21** | 2200.10331 | L | **423.2602** | **3** |
| **22** | 2363.16664 | Y | **310.17613** | **2** |
| **23** | | K | 147.1128 | **1** |

**Table S3.** 234 Aha sfGFP mass spec expected fragmentation for example peptide

| | b+ | Seq. | $y^+$ | |
|---|---|---|---|---|
| **1** | 116.03422 | D | | **23** |
| **2** | **253.09313** | H | 2410.25137 | **22** |
| **3** | **324.13025** | A | 2273.19246 | **21** |
| **4** | **423.19866** | V | 2202.15534 | **20** |
| **5** | **536.28272** | L | 2103.08693 | **19** |
| **6** | **649.36679** | L | 1990.00287 | **18** |
| **7** | **778.40938** | E | 1876.9188 | **17** |
| **8** | **925.47779** | F | 1747.87621 | **16** |
| **9** | **1024.54621** | V | 1600.80779 | **15** |
| **10** | 1125.59389 | T | 1501.73938 | **14** |
| **11** | 1196.631 | A | **1400.6917** | **13** |
| **12** | **1267.66811** | A | **1329.65459** | **12** |
| **13** | 1324.68958 | G | **1258.61747** | **11** |
| **14** | 1437.77364 | I | **1201.59601** | **10** |
| **15** | 1538.82132 | T | **1088.51195** | **9** |
| **16** | 1675.88023 | H | **987.46427** | **8** |
| **17** | 1732.9017 | G | **850.40536** | **7** |
| **18** | 1858.95586 | M-Met->Aha | 793.38389 | **6** |
| **19** | 1973.9828 | D | 667.32973 | **5** |
| **20** | 2103.02539 | E | 552.30279 | **4** |
| **21** | 2216.10946 | L | 423.2602 | **3** |
| **22** | 2379.17278 | Y | **310.17613** | **2** |
| **23** | | K | 147.1128 | **1** |

**Table S4.** 150 Aha sfGFP mass spec expected fragmentation for example peptide

| | **b⁺** | **Seq.** | **y⁺** | |
|---|---|---|---|---|
| **1** | 114.09134 | L | | 16 |
| **2** | 243.13393 | E | 1826.84564 | 15 |
| **3** | 406.19726 | Y | 1697.80304 | 14 |
| **4** | 520.24019 | N | 1534.73971 | 13 |
| **5** | 667.3086 | F | 1420.69679 | 12 |
| **6** | 781.35153 | N | 1273.62837 | 11 |
| **7** | 868.38356 | S | 1159.58545 | 10 |
| **8** | 1005.44247 | H | 1072.55342 | 9 |
| **9** | 1131.49663 | M-Met->Aha | 935.49451 | 8 |
| **10** | 1230.56505 | V | 809.44035 | 7 |
| **11** | 1393.62837 | Y | 710.37193 | 6 |
| **12** | 1506.71244 | I | 547.3086 | 5 |
| **13** | 1607.76012 | T | 434.22454 | 4 |
| **14** | 1678.79723 | A | 333.17686 | 3 |
| **15** | 1793.82417 | D | 262.13975 | 2 |
| **16** | | K | 147.1128 | 1 |

**Table S5**. 150 Cha sfGFP mass spec expected fragmentation for example peptide

| | $b^+$ | Seq. | $y^+$ | |
|---|---|---|---|---|
| **1** | 114.09134 | L | | |
| **2** | 243.13393 | E | 1810.83949 | **15** |
| **3** | 406.19726 | Y | 1681.7969 | **14** |
| **4** | 520.24019 | N | 1518.73357 | **13** |
| **5** | 667.3086 | F | 1404.69064 | **12** |
| **6** | 781.35153 | N | 1257.62223 | **11** |
| **7** | 868.38356 | S | 1143.5793 | **10** |
| **8** | 1005.44247 | H | 1056.54727 | **9** |
| **9** | 1115.49048 | M-Met->Cha | 919.48836 | **8** |
| **10** | 1214.5589 | V | 809.44035 | **7** |
| **11** | 1377.62223 | Y | 710.37193 | **6** |
| **12** | 1490.70629 | I | 547.3086 | **5** |
| **13** | 1591.75397 | T | 434.22454 | **4** |
| **14** | 1662.79108 | A | 333.17686 | **3** |
| **15** | 1777.81802 | D | 262.13975 | **2** |
| **16** | | K | 147.1128 | **1** |

**Table S6**. 150 oxidized Cha sfGFP mass spec expected fragmentation for example peptide

| | $b^+$ | Seq. | $y^+$ | |
|---|---|---|---|---|
| **1** | 114.09134 | L | | |
| **2** | 243.13393 | E | 1826.83898 | **15** |
| **3** | 406.19726 | Y | 1697.79639 | **14** |
| **4** | 520.24019 | N | 1534.73306 | **13** |
| **5** | 667.3086 | F | 1420.69013 | **12** |
| **6** | 781.35153 | N | 1273.62172 | **11** |
| **7** | 868.38356 | S | 1159.57879 | **10** |
| **8** | 1005.44247 | H | 1072.54676 | **9** |
| **9** | 1131.48997 | M-Met->OxCha | 935.48785 | **8** |
| **10** | 1230.55839 | V | 809.44035 | **7** |
| **11** | 1393.62172 | Y | 710.37193 | **6** |
| **12** | 1490.70629 | I | 547.3086 | **5** |
| **13** | 1506.70578 | T | 434.22454 | **4** |
| **14** | 1662.79108 | A | 333.17686 | **3** |
| **15** | 1678.79057 | D | 262.13975 | **2** |
| **16** | | K | 147.1128 | **1** |

**Table S7 is a separate Excel attachment**

**Table S8.** Melting temperatures of each sfGFP variant melt

| Protein | Tm (°C) | Error |
|---|---|---|
| **150 Met** | 80.5 | 0.1 |
| | 79.3 | 0.2 |
| | 78.9 | 0.2 |
| **150 Aha** | 81.6 | 0.3 |
| | 81.2 | 0.1 |
| | 80.1 | 0.3 |
| **150 Cha** | 78.2 | 0.2 |
| | 79.0 | 0.2 |
| | 80.1 | 0.5 |
| **234 Met** | 79.3 | 0.1 |
| | 78.8 | 0.1 |
| | 80.2 | 0.1 |
| **234 Aha** | 78.9 | 0.2 |
| | 75.2 | 0.3 |
| | 78.5 | 0.1 |
| **234 Cha** | 73.0 | 0.3 |
| | 77.0 | 0.2 |
| | 78.2 | 0.3 |

**Table S9.** sfGFP variant genes

| | |
|---|---|
| 150<br>sfGFP | ATGGCGAGCAAAGGCGAGGAACTGTTCACCGGTGTGGTTCCGATCCTG<br>GTGGAGCTGGACGGCGATGTTAACGGTCACAAGTTTAGCGTGCGTGGT<br>GAGGGCGAAGGTGACGCGACCAACGGCAAGCTGACCCTGAAATTCATT<br>TGCACCACCGGTAAACTGCCGGTGCCGTGGCCGACCCTGGTTACCACC<br>CTGACCTACGGCGTGCAGTGCTTTAGCCGTTATCCGGACCACATCAAGC<br>GTCACGATTTCTTTAAAAGCGCGCTGCCGGAGGGCTACGTTCAAGAAC<br>GTACCATTAGCTTCAAGGACGATGGTACCTATAAAACCCGTGCGGAAGT<br>GAAGTTTGAAGGCGACACCCTGGTTAACCGTATCGAGCTGAAGGGTATT<br>GACTTCAAAGAAGATGGCAACATCCTGGGTCACAAGCTGGAGTACAAC<br>TTTAACAGCCACATG<br>GTTTATATTACCGCGGATAAGCAGAAAAACGGCATCAAGGCGAACTTTA<br>AAATTCGTCACAACGTGGAAGACGGTAGCGTTCAACTGGCGGATCACT<br>ACCAGCAAAACACCCCGATTGGTGATGGTCCGGTGCTGCTGCCGGATAA<br>CCACTATCTGAGCACCCAGAGCGTTCTGAGCAAGGACCCGAACGAGAA<br>ACGTGATCACGCGGTGCTGCTGGAATTCGTTACCGCGGCGGGTATTACC<br>CATGGTGCGGATGAACTGTACAAAGGTAGCCACCACCACCACCACCAC<br>TAA |
| 234<br>sfGFP | ATGGCGAGCAAAGGCGAGGAACTGTTCACCGGTGTGGTTCCGATCCTG<br>GTGGAGCTGGACGGCGATGTTAACGGTCACAAGTTTAGCGTGCGTGGT<br>GAGGGCGAAGGTGACGCGACCAACGGCAAGCTGACCCTGAAATTCATT<br>TGCACCACCGGTAAACTGCCGGTGCCGTGGCCGACCCTGGTTACCACC<br>CTGACCTACGGCGTGCAGTGCTTTAGCCGTTATCCGGACCACATCAAGC<br>GTCACGATTTCTTTAAAAGCGCGCTGCCGGAGGGCTACGTTCAAGAAC<br>GTACCATTAGCTTCAAGGACGATGGTACCTATAAAACCCGTGCGGAAGT<br>GAAGTTTGAAGGCGACACCCTGGTTAACCGTATCGAGCTGAAGGGTATT<br>GACTTCAAAGAAGATGGCAACATCCTGGGTCACAAGCTGGAGTACAAC<br>TTTAACAGCCACAAC<br>GTTTATATTACCGCGGATAAGCAGAAAAACGGCATCAAGGCGAACTTTA<br>AAATTCGTCACAACGTGGAAGACGGTAGCGTTCAACTGGCGGATCACT<br>ACCAGCAAAACACCCCGATTGGTGATGGTCCGGTGCTGCTGCCGGATAA<br>CCACTATCTGAGCACCCAGAGCGTTCTGAGCAAGGACCCGAACGAGAA<br>ACGTGATCACGCGGTGCTGCTGGAATTCGTTACCGCGGCGGGTATTACC<br>CATGGTATGGATGAACTGTACAAAGGTAGCCACCACCACCACCACCAC |
